# Volumetric imaging and computation to explore contractile function in zebrafish hearts

**DOI:** 10.1101/2024.11.14.623621

**Authors:** Alireza Saberigarakani, Riya P. Patel, Milad Almasian, Xinyuan Zhang, Jonathan Brewer, Sohail S. Hassan, Jichen Chai, Juhyun Lee, Baowei Fei, Jie Yuan, Kelli Carroll, Yichen Ding

**Affiliations:** Department of Bioengineering, The University of Texas at Dallas, Richardson, TX 75080, USA; Department of Biology, The University of Texas at Dallas, Richardson, TX 75080, USA; Department of Bioengineering, The University of Texas at Arlington, Arlington, TX 76019, USA; Center for Imaging and Surgical Innovation, The University of Texas at Dallas, Richardson, TX 75080, USA; Department of Biology, Wofford College, Spartanburg, SC 29303, USA; Hamon Center for Regenerative Science and Medicine, UT Southwestern Medical Center, Dallas, TX 75390, USA

**Keywords:** Light field microscopy, virtual reality, cell tracking, cardiac contractility

## Abstract

Despite advancements in cardiovascular engineering, heart diseases remain a leading cause of mortality. The limited understanding of the underlying mechanisms of cardiac dysfunction at the cellular level restricts the development of effective screening and therapeutic methods. To address this, we have developed a framework that incorporates light field detection and individual cell tracking to capture real-time volumetric data in zebrafish hearts, which share structural and electrical similarities with the human heart and generate 120 to 180 beats per minute. Our results indicate that the in-house system achieves an acquisition speed of 200 volumes per second, with resolutions of up to 5.02 ± 0.54 µm laterally and 9.02 ± 1.11 µm axially across the entire depth, using the estimated-maximized-smoothed deconvolution method. The subsequent deep learning-based cell trackers enable further investigation of contractile dynamics, including cellular displacement and velocity, followed by volumetric tracking of specific cells of interest from end-systole to end-diastole in an interactive environment. Collectively, our strategy facilitates real-time volumetric imaging and assessment of contractile dynamics across the entire ventricle at the cellular resolution over multiple cycles, providing significant potential for exploring intercellular interactions in both health and disease.

## Introduction

Heart diseases are a leading cause of death worldwide, responsible for 31% of all fatalities ^1,2^. The development of targeted therapies and interventions depends on a deep understanding of intercellular interactions throughout the heart. Although significant progress has been made, effective methods to study cardiac contractile function at the cellular level in a beating heart are still needed. Recent advancements have shown that zebrafish are an excellent model for heart studies due to their genetic flexibility, optical transparency, and similarities in heart structure and electrocardiograms to humans ^3–5^. This makes them a valuable tool for investigating cardiac structure and function in a living organism.

While the imaging power of zebrafish has been established ^3–5^, technical limitations remain to be fully addressed. Widefield microscopy allows for the rapid acquisition of contracting heart in zebrafish larvae, but it comes at the cost of limited spatial resolution and out-of-focus background. On the other hand, confocal and multiphoton microscopy provide superb spatial resolution and three-dimensional (3D) information. Still, the reduced imaging speed and increased phototoxicity compromise their utility for *in vivo* volumetric studies of cardiac contraction in zebrafish ^6–8^. While recent methods such as light-sheet microscopy along with synchronization ^9–16^ have enabled the decent 4D (3D spatial + 1D temporal) imaging of zebrafish heart, the inherent capability of capturing a cross-sectional image at a single time point constrains its volumetric acquisition at a pace of 2 to 3 beats per second ^15–17^, regardless of arrhythmic heartbeats. In this context, we sought to leverage the emerging light field microscopy (LFM) ^18–22^ to assess cardiac structure and contractile function in zebrafish larvae.

In this study, we developed a framework ranging from the LFM imaging system to interactive tracking of individual cells, using transgenic zebrafish labeling myosin light chain (*Tg(myl7:nucGFP)*) as our model system. To capture instantaneous dynamics of cardiomyocytes in the contracting heart with minimal autofluorescence background and optimal orientation, we integrated selective volumetric illumination with LFM detection, rather than using the bright-field illumination in the inverted microscopes typically applied for zebrafish studies ^22,23^. For 3D volumetric reconstruction from a 2D snapshot, we assessed two representative deconvolution methods and quantified their performance on the cardiac images, bypassing the large training datasets in deep learning methods ^24,25^. We further utilized 3DeeCellTracker ^26^ and our interactive virtual reality (VR) method ^27–30^ for individual cell tracking across the intact contracting heart. Collectively, this strategy enables real-time volumetric investigation of 4D time-dependent cardiac contraction with cellular resolution from ventricular end-systole to end-diastole, holding great promise for the advanced understanding of intercellular interaction under physiological and pathophysiological conditions.

## Results

### Fundamental concept of LFM and system implementation

LFM introduces a new dimension by modulating the native image plane to capture the angular information from the specimen, rather than detecting the intensity and position of fluorescence emitters on the focal plane. We introduced a microlens array (MLA) on the native image plane to project incoming angular views to corresponding pixels behind each lenslet ^24^. To illustrate the mechanism of angular view detection, on-axis (red) and off-axis rays (blue) passing through a specific lenslet converge to different pixels on the detector (**Fig. 1A**). This suggests that identical pixels behind the lenslets correspond to a unique angular view. By extracting the identical pixels and concatenating them in the order of their original lenslets, the specific angular perspectives from the specimen were captured (color-coded in **Fig. 1B**). Due to the ability of LFM to collect different angular views, varying depths of the point emitter result in distinct patterns in the collected data, which can be decoded to extract depth-related information. When the emitter is positive defocus, it illuminates the pixels leaning toward the center lenslet in each neighbor lenslet (**Fig. 1C**). Conversely, moving it further away illuminates the opposite side of the lenslets compared to the center lenslet. Thus, the asymmetrical point spread function (PSF) along the Z-axis leads to the extraction of the 3D depth information from 2D images (**Fig. 1D**). To reduce the autofluorescence of the yolk sac in zebrafish larvae and enhance the image contrast in cardiac imaging, we selectively illuminated the zebrafish heart by a rod-shaped laser beam ^25,31^. Due to the density of fluorescence distribution in the transgenic models, we utilized both Richardson-Lucy (RL) and Estimate-Maximize-Smooth (EMS) ^32^ deconvolution methods to reconstruct the 4D dynamics from time-dependent 2D snapshots, and evaluated the results of these two methods qualitatively and quantitatively.

**Figure 1.**
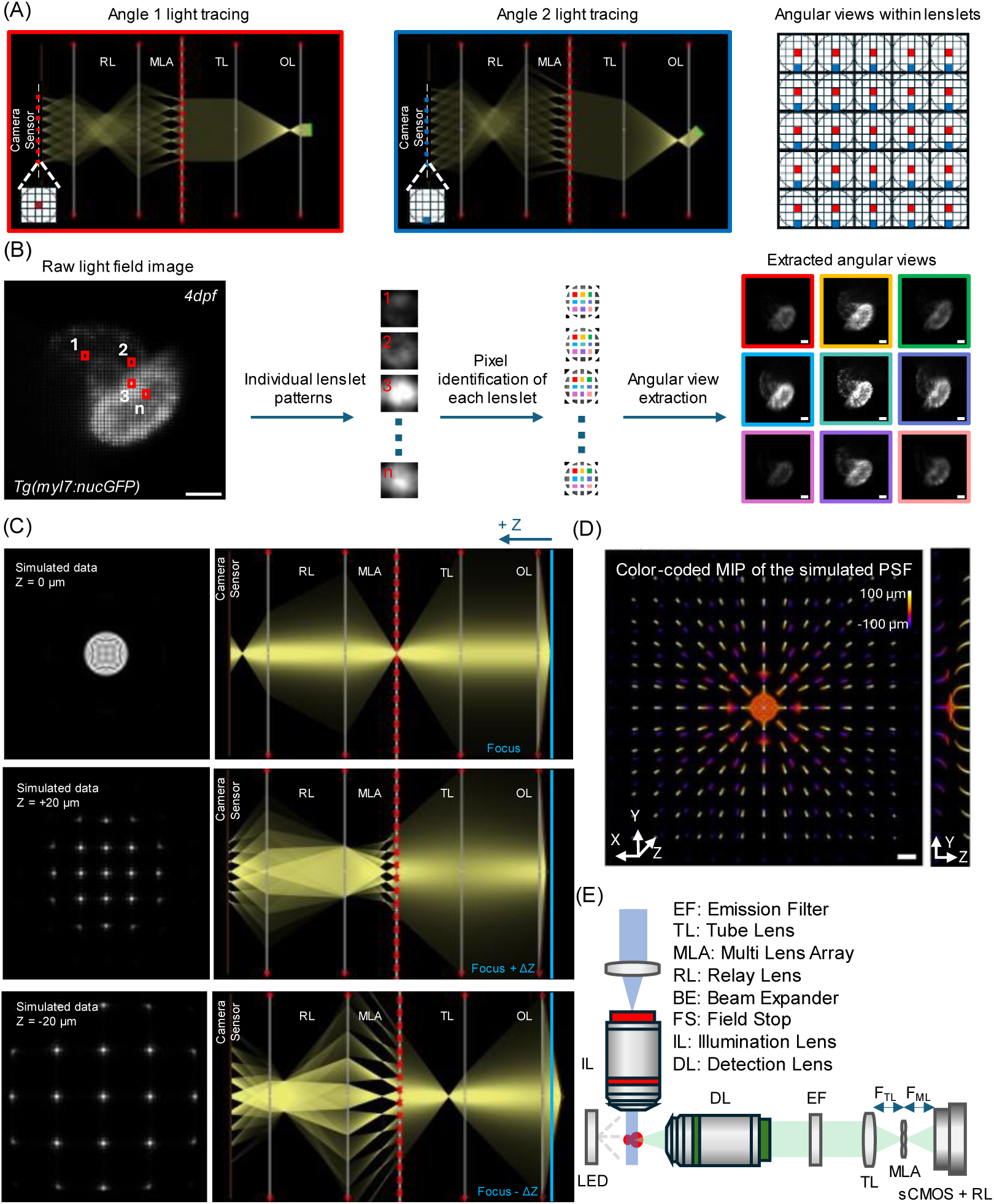
Light field microscopy (LFM) configuration and system construction. (A) Distinctive patterns of different angular information (red and blue) on the sCMOS sensor behind the lenslets. (B) Process of extracting different angular views from a zebrafish heart, including pixel identification in the captured image and reordering them to create multiple low-resolution views of the heart. (C) Simulation of the pattern on the detector from a point emitter at varying axial positions. (D) Maximum intensity projection (MIP) of the PSF in both lateral and axial views. (E) Schematic of the in-house LFM with selective volume side illumination. Scale bars: 50 µm.

### Reconstruction process and PSF calibration

The computational LFM image reconstruction process involves two main steps. The first step is rectification, where we define the location of each lenslet and the pixels behind them in the captured data. In our rectangular lenslet arrangement, we assigned an area of 15×15 pixels to each lenslet. In the second step, we applied 15 iterations of deconvolution using the theoretical PSF to obtain the final reconstructed volume corresponding to each timestep of the imaging process (**Fig. 2A**).

**Figure 2.**
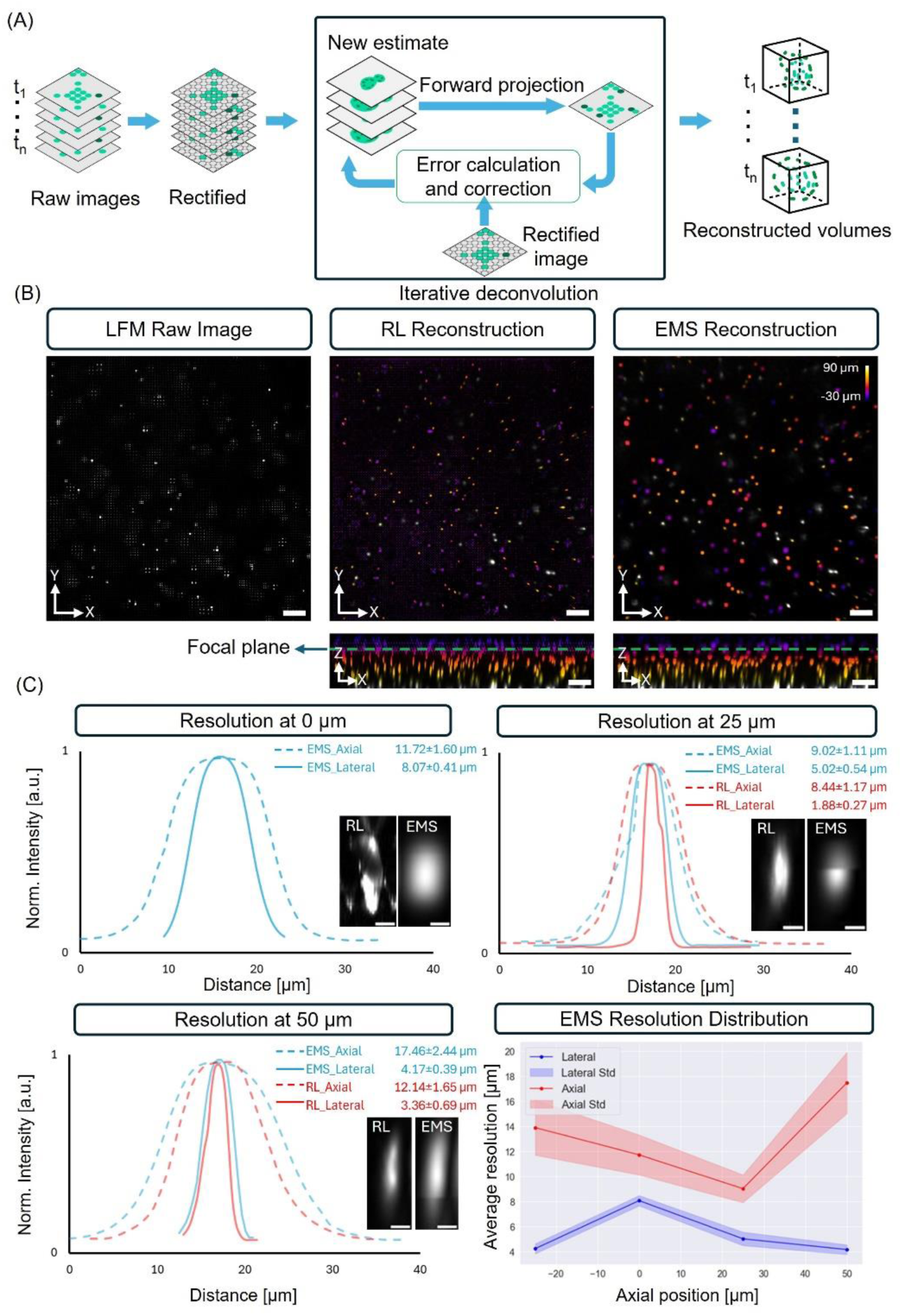
Reconstruction process and PSF calibration. (A) Schematic of the 4D reconstruction process, including image rectification and pixel identification, iterative deconvolution for each timestep to reconstruct a 3D volume from 2D input, and assembling the 3D reconstructed files to generate the 4D data. (B) MIP reconstruction of fluorescent beads using EMS and RL in both axial and lateral views, with color-coded depth information. Scale bar: 50 µm. (C) Lateral and axial resolution from both deconvolution methods at different depths, averaging measurements over 10 beads under each condition. (D) Distribution of lateral and axial resolution across varying depths in EMS reconstruction. Scale bar: 5 µm.

To quantify the resolution of this custom-made LFM system, we imaged fluorescent beads with a diameter of 0.53 µm at a concentration of 1:10^5^. We reconstructed the raw image over a 120 µm depth, ranging from a negative defocus of 30 µm to positive defocus of 90 µm beyond the focal plane. After rectification, the representative reconstruction of maximum intensity projection (MIP) images using both RL and EMS methods along XY and XZ planes were presented (**Fig. 2B**). Each method demonstrated varying axial and lateral resolutions at different depths. We measured the representative full width at half maximum (FWHM) at depths of 0, 25, and 50 µm from the focal plane. Our results indicated that artifacts significantly dominate at the focus in RL method, while EMS improved the reconstruction quality on the focal plane but showed artifacts across the 3D volume (**Fig. 2C** and **Supplementary Fig. S1**). The quantitative results of the lateral (solid) and axial (dashed) resolutions at representative depths of 0, 25, and 50 µm using EMS were 8.07±0.41 and 11.72±1.60 µm, 5.02±0.54 and 9.02±1.11 µm, and 4.17±0.39 and 17.46±2.44 µm, respectively, based on an average of 10 bead measurements for each condition. In contrast, the resolutions of RL method at the depths of 25 and 50 µm were 1.88±0.27 and 8.44±1.17 µm, and 3.36±0.69 and 12.14±1.65 µm, respectively. The results indicate that RL provides better lateral and axial resolution compared to EMS. However, the image quality around the focal plane is significantly degraded due to strong artifacts, consistent with other reports ^19,32,33^. In contrast, EMS produces milder artifacts and more consistent resolutions under the same condition, making it a more suitable choice for deconvolution-based LFM when uniformity across different depths is essential. Further analysis reveals that the optimal resolution in EMS occurs approximately 25 µm closer to the objective lens relative to the focal plane (**Fig. 2D**), establishing the optimal range for enhanced imaging clarity in subsequent experiments.

### Volumetric imaging of the intact heart in zebrafish larvae

To ensure the zebrafish maintained the correct orientation during data acquisition, we mounted the larva in an FEP tube using 1% agarose after soaking it in a 150 mg/L MS-222 (Tricaine) solution for 10 minutes. We focused on the detection objective lens closer to the ventral side of the ventricular myocardium to ensure the high contrast. This setup allowed us to exclude most of the yolk while keeping a clear view of the entire heart (**Fig. 3A**). We imaged both the ventricle and atrium of transgenic *Tg(myl7:nucGFP)* zebrafish larvae from 3 to 5 days post-fertilization (dpf) with the exposure time of 5 ms to investigate myocardial structure and contractile function across the entire heart, resulting in the recording of 200 volumes per second. After 15 iterations at each timestep, the heart reconstruction could be rendered as a 3D model or illustrated using MIP images in both lateral and axial views, with cellular resolution (**Fig. 3B**). To improve the efficiency of analyses, such as heart rate in the reconstruction, we developed a custom algorithm to automatically track the myocardial boundary in a 2D plane along the radial axis. By tracking cellular movements from systole to diastole, we can map the contraction-relaxation patterns at various depths over three cardiac cycles, providing a basis for computationally tracking the instantaneous displacement and velocity of individual cells (**Fig. 3C** and **Supplementary Video S1**).

**Figure 3.**
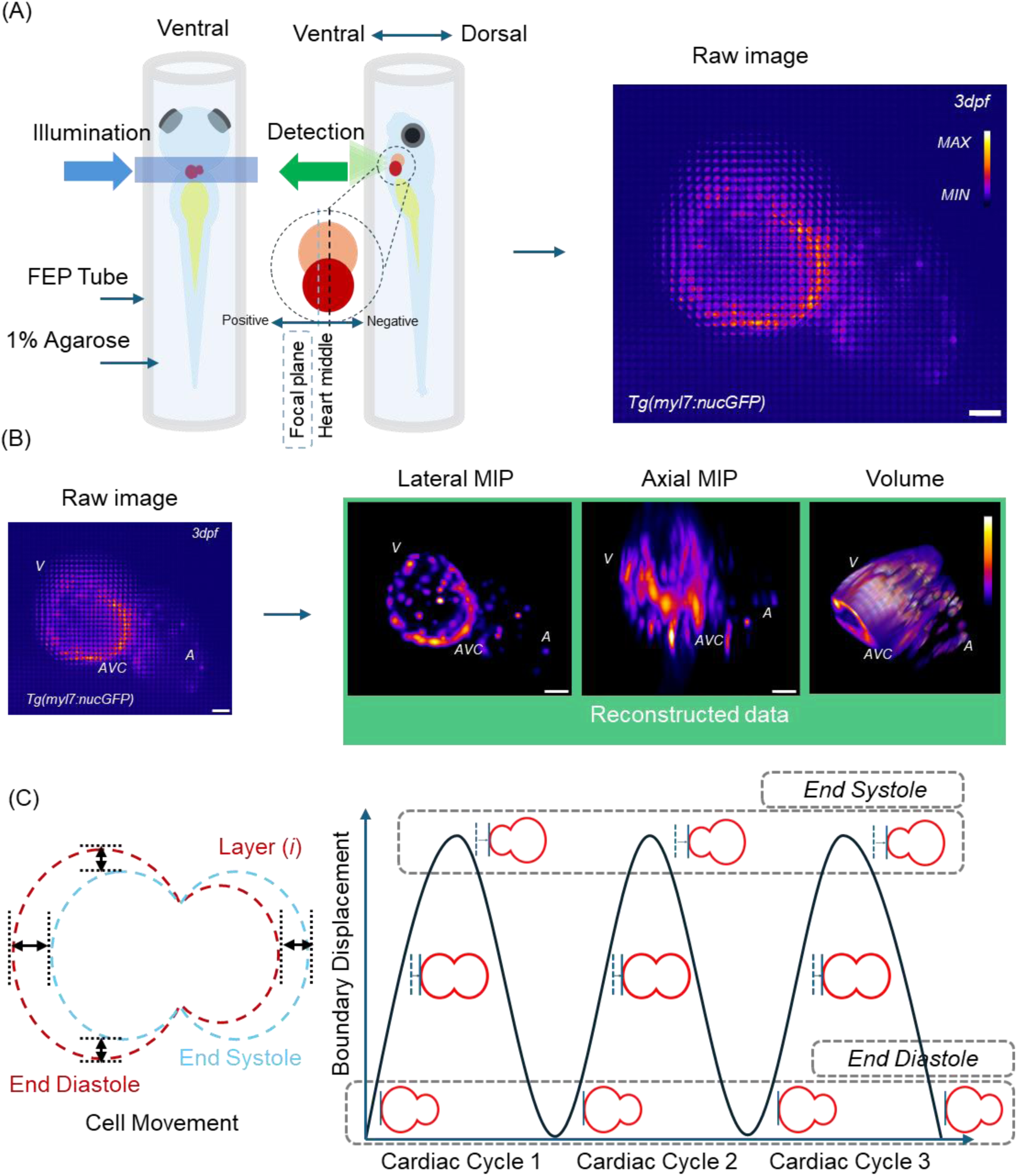
Zebrafish mounting, reconstruction results, and heartbeat analysis procedure. (A) Schematic of the zebrafish mounting procedure in agarose and FEP tube with proper orientation to capture data from both chambers. (B) Acquired data after reconstruction of each timestep, including MIP in the lateral view and the 3D volume of the region of interest. A: atrium; V: ventricle; AVC: atrioventricular canal. (C) Framework illustration for extracting valuable data such as heartbeat, cyclic cellular patterns, initiation delays, cellular movement, and speed from each layer of the reconstructed 4D data. Scale bar: 50 µm.

### Reconstruction of 4D contracting heart

Using either the RL or EMS method, we reconstructed the zebrafish heart from 3 dpf to 5 dpf and assessed the results qualitatively and quantitatively (**Fig. 4**). While both methods provide detailed 3D volumes of the heart at different depths, EMS, with its anti-aliasing component, reduces artifacts in layers near the focal plane, resulting in smoother image quality in reconstructed layers. This is evident in the MIP of the whole reconstructed volumes and is consistent with the calibration results from fluorescent beads (**Fig. 2**). To investigate the differences in cell reconstruction quality between two methods, we examined the intensity distribution across the cells and found that near the focal plane, EMS demonstrates a more consistent Gaussian distribution, while RL predominantly shows spike artifacts, making cell identification more challenging in both individual slice and MIP images. The intensity profiles of representative cells at 3 and 4 dpf are presented in green. To assess the capability of both methods in preserving different frequency components in the reconstructed volumes, we further performed a Fourier transform on the MIP results. The analysis revealed more high-frequency components in the spectrum of the RL method, contributing to the degradation of the reconstructed image quality. In contrast, the EMS method preserves the structural integrity of the myocardium while minimizing artifacts. To enable high-throughput reconstruction of cardiac contraction over the time course, we further implemented parallel computing in EMS in this study ^19, 32^.

**Figure 4.**
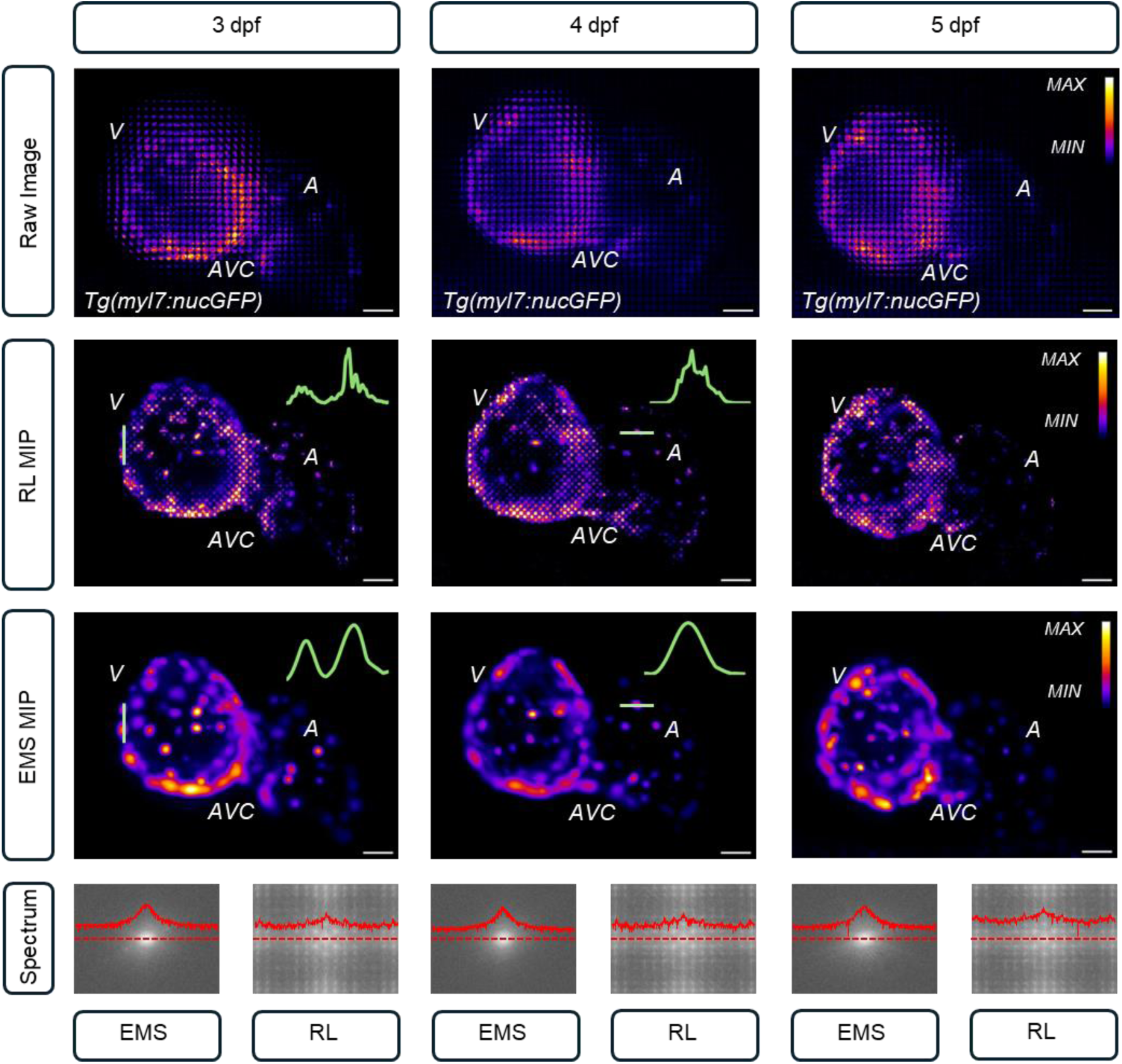
RL and EMS reconstruction of zebrafish hearts in spatial and frequency domains. MIP images, along with the highlighted green line profile in zebrafish larvae, indicate that EMS resolves cells with reduced artifacts across the entire heart, whereas RL exhibits dominant artifacts around the focal plane. The logarithmic Fourier transform of the images illustrates the presence of high frequency components and their impact on obscuring primary signals. A: atrium; V: ventricle; AVC: atrioventricular canal. Scale bar: 25 µm.

### LFM-assisted tracking of contractile function with cellular resolution

LFM combined with the individual cell tracking allows for the assessment of the global cardiac contraction from end-systole to end-diastole, as well as the identification of differences in representative cross-sections of the zebrafish heart across five cardiac cycles. After ten hours of parallel computation, we reconstructed 15 volumes of the zebrafish heart at 3 dpf using 15 iterations of EMS deconvolution, resulting in a complete reconstruction of the entire cardiac cycle in approximately ninety hours. To investigate myocardial contraction and relaxation with cellular resolution, we demonstrated representative cross-sections at depths of -20, 0, and 20 µm relative to the focal plane, and created depth-color-coded MIP images for each timestep (**Fig. 5**), illustrating the cardiac contractility within 80 µm range (**Supplementary Video S2**). We tracked the myocardial boundary highlighted in orange at various depths to analyze regional contracting phases, displacement, and velocity throughout the complete cardiac cycle (**Fig. 5A**). Using the end-diastole as the baseline for all layers, our results unravel regional differences in displacement and velocity across the entire ventricle, with a 30% increase in radial displacement (24 to 36.4 µm) and velocity (0.2 to 0.31 µm/ms) observed in the coronal plane from the central section of the heart toward the epicardium. By analyzing the starting and ending points of each cardiac cycle across varying depths, we observed an approximate 40 ms initiation delay as we moved towards the ventral wall of the heart. To further investigate, we examined another zebrafish at 6 dpf and found a similar delay in the initiation of cardiac pacing between central and peripheral heart layers, with an approximate 45 ms delay corresponding to a 40 µm depth difference (**Fig. 5B**). This suggests that rapid LFM data acquisition holds great promise for investigating mechanical contractile dysfunction from pacing cells to the apex across the intact heart.

**Figure 5.**
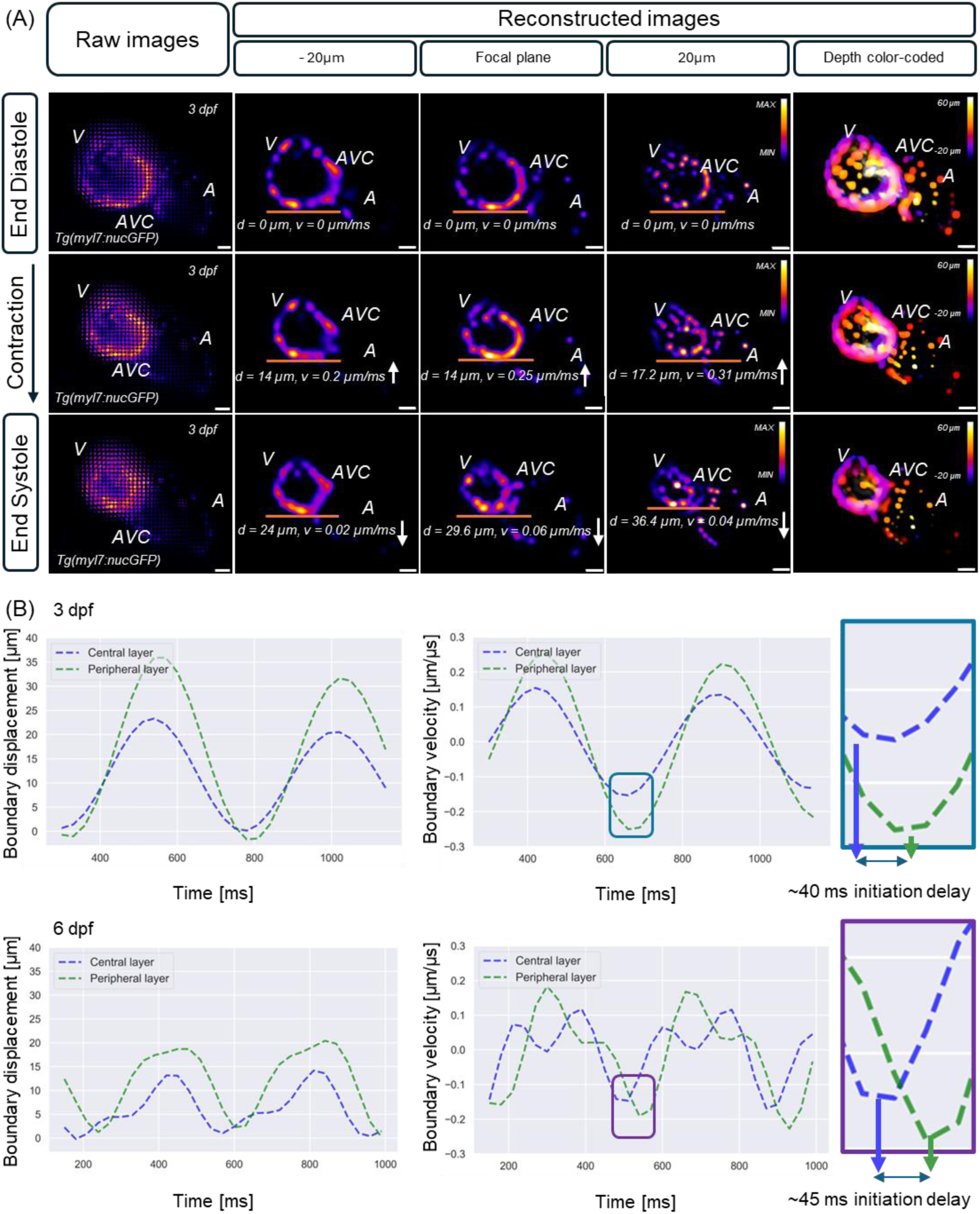
Analysis of cardiac contractile function in zebrafish larvae. (A) Instantaneous displacement and velocity of individual layers at depths of -20, 0, and 20 µm relative to the focal plane are quantified at different cardiac phases over five cycles in the zebrafish larva at 3 dpf. Depth-color-coded MIP projections illustrate the global changes across varying depths at each timestep. Pseudo-color indicates the location of cells across different layers in the depth-color-coded results. (B) Displacement and velocity patterns in the ventricle illustrate the initiation delay of contraction cycles at varying layers in zebrafish larvae at 3 and 6 dpf, reflecting the sequential mechanical contraction driven by electrical conduction from pacing cells in the atrium to the ventricular apex. Scale bar: 25 µm.

To explore region-specific heterogeneity in cellular activity patterns, we implemented 3DeeCellTracker ^26^ to automatically segment and track individual cardiomyocytes. We quantified the displacement and velocity of two fully resolved cells during consecutive cardiac cycles along X, Y, and Z axes. Specifically, the maximum magnitudes of displacement for cell #1 and #2, located on the ventricular wall, are 90 µm and 52 µm, respectively, with corresponding speeds of 0.55 µm/ms and 0.21 µm/ms in two successive cycles, suggesting varying contractility and a potential heterogenous distribution of myocardial properties across the entire ventricle (**Fig. 6A**). Given the presence of numerous traveling cells throughout the heart, we further demonstrated the feasibility of our customized VR environment for effective data handling and interpretation using segmented cells from each timestep. The interactive and manipulative nature of this method allows us to investigate specific cells of interest in arbitrary regions and study their displacement and velocity, providing an entry point for biomedical professionals with limited engineering training to assess intracellular interactions within 4D complicated structures and functional dynamics using intuitive functions. We utilized a color-coded velocity map to analyze variations in cell velocity across different heart regions. In this map, lighter colors represent higher speeds and greater displacements than other cells. Additionally, the graphical user interface provides information on the relative distances among selected cells, allowing for the study of cell-cell kinematic interactions during multiple cardiac contractions (**Fig. 6B** and **Supplementary Video S3**).

**Figure 6.**
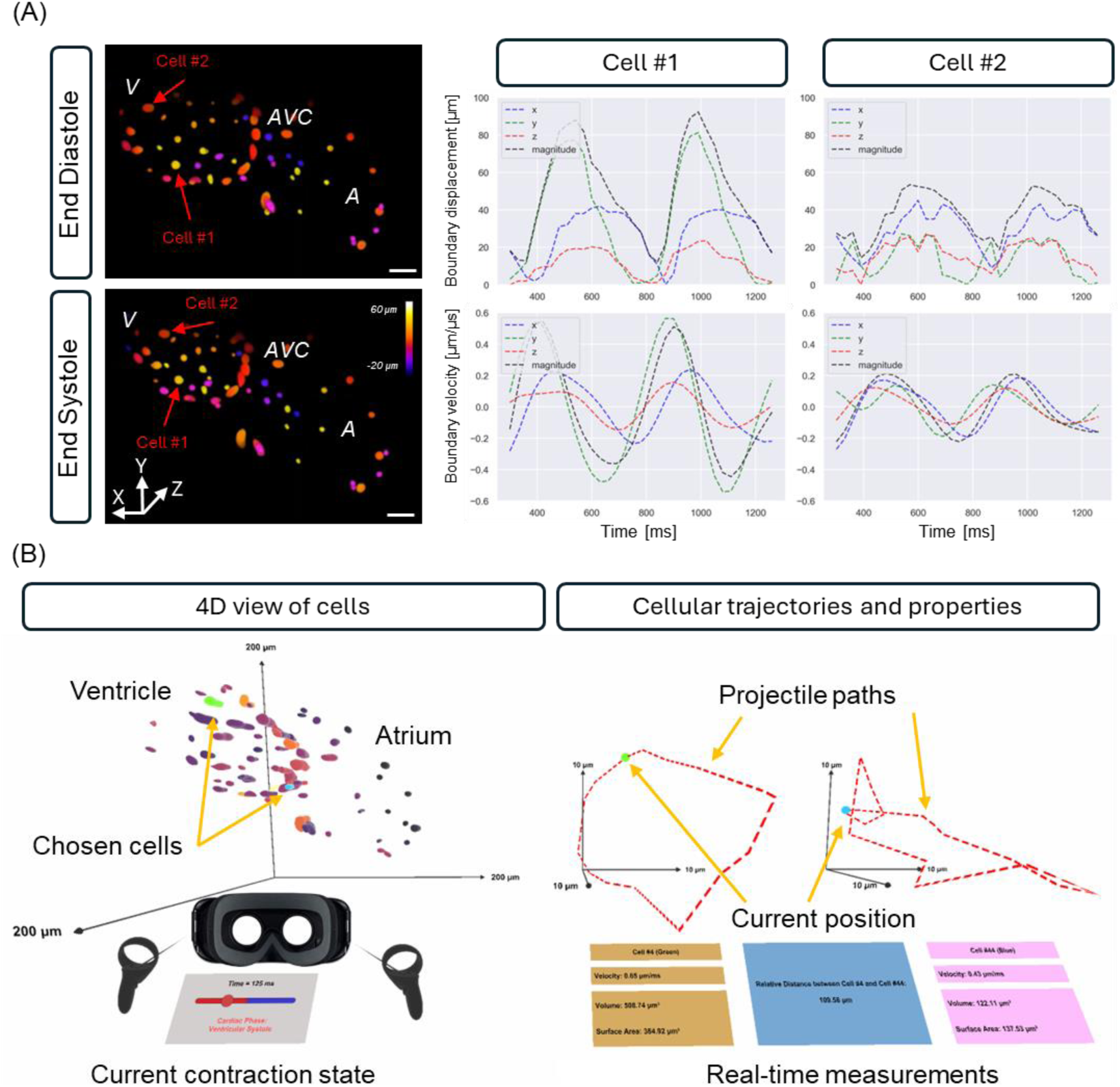
Cell tracking results in zebrafish beating heart in standard and virtual reality modes. (A) The 3DeeCellTracker enables us to segment and track fully resolved individual cells over two consecutive cardiac cycles. Displacement and velocity patterns of Cells #1 and #2 located in the ventricle depict region-specific contractility across the cycle. (B) Graphical user interface displaying a virtual representation of cardiac contractions. The interface dynamically provides the trajectory paths, speeds, and relative distance between two cells, as well as the surface area and volume of the selected cells.

## Discussion

In this study, we developed a framework to uncover the 4D pattern of cardiac contractile function with cellular resolution in zebrafish larvae. First, we built an in-house LFM imaging system to capture the intact beating zebrafish heart, obviating the mechanical slicing and synchronization of varying cross-sections. We employed both RL and EMS deconvolution methods to reconstruct the 3D cardiac volume from captured 2D snapshots. Our results indicated that RL provided higher resolution further from the focal plane, while EMS reduced artifacts closer to the focal plane in cardiac images. Further comparison of the resolution at various depths suggests that RL provides overall better resolution with the same number of iterations, compared to the EMS method. This is supported by quantitative analysis at a depth of 25 µm, showing a 60% improvement in lateral resolution and a 5% improvement in axial resolution, respectively. However, due to the limited depth resolution in the axial view with LFM, EMS is considered a more effective approach for reconstructing zebrafish heart data, as it reduces artifacts and enhances structural clarity. Our findings reveal significant regional variations in both the displacement and velocity of individual cardiomyocytes within the ventricle. Specifically, we observed up to a 30% difference between cardiomyocytes located in the widest 2D cross-sectional layers along the longitudinal axis of the ventricle, compared to those positioned closer to the ventral (anterior) wall of the heart. In addition, the initiation delay among different layers in the reconstructed heart aligns with the physiological process of myocardial contraction, highlighting the potential application of this method for exploring electrical conduction and mechanical contraction from the sinoatrial node to the apex.

Subsequently, we applied the 3DeeCelltracker algorithm ^26^ to segment all fully resolved cells, enabling us to track individual cells in instantaneous cardiac dynamics with cellular resolution. Due to the varying spatial resolution at different depths in LFM, manual annotation is required to validate whether all cells of interest are accurately tracked. Our current representative cell movement patterns allow for identifying variations in cellular displacement, velocity, and pace across different regions of the ventricle, from the center to the ventral wall. Further studies using this method hold great promise for uncovering myocardial properties and mechanical stress across different regions of the intact heart, as well as exploring the interaction between hemodynamics and cardiac contraction during heart development and regeneration. Collectively, our method provides a new strategy for *in vivo* study of 4D cardiac contractile function with cellular resolution in zebrafish larvae, enabling us to investigate instantaneous cardiac dynamics in beating hearts. Unlike retrospective synchronization used in light-sheet imaging ^12,14^, this framework offers a pathway for investigating hearts with abnormalities and contractile dysfunctions during active contraction, holding the great potential to uncover the mechanism underlying cardiac morphogenesis and facilitate new therapies.

### Limitations

While we were able to capture and track individual cardiomyocytes in the 4D contracting heart, the high temporal resolution of LFM is achieved at the cost of limited axial resolution, resulting in varying resolving power for cell tracking at different depths. Our results suggest that an in-depth analysis of complete ventricular, rather than atrial, contraction can be effectively conducted in zebrafish larvae from 3 to 6 dpf. Our previous study demonstrates that the acquisition speed of up to 200 volumes per second is required for the capture of 2-3 cardiac cycles in 4D without aliasing ^33^, and therefore, the trade-off between imaging speed, fluorescence photon budget, and reconstruction efficiency remains to be fully determined. Our current results indicate that the exposure time of 5 ms and parallel computation-assisted deconvolution enable to resolve the 4D contracting heart in zebrafish. Additionally, the variation of fluorescence intensity in different zebrafish larvae also poses a challenge to reproduce the experiments under the same technical conditions. Our results suggest that high-throughput screening is still essential when conducting LFM experiments on the same batch of zebrafish samples within a narrow developmental window.

With regard to the tracking of a subset of cells, the primary reason is that 3DeeCellTracker demonstrates higher accuracy in the XY plane compared to the Z-axis. In our study, the varying axial resolution within the 3D volume (ranging from 8 to 17 µm) complicates consistent tracking, particularly for cells that stretch more than others. Additionally, while LFM reconstruction excels with sparse signals, when the heart chambers enter the systole phase, cells become densely packed, posing a significant challenge to differentiate and track them along both lateral and axial directions in the following image post-processing. To address these limitations, improving the spatial resolution of the imaging system and reconstruction accuracy with the advanced algorithms for dense signals ^34^ are both essential. Recent advances in deep learning have also allowed us to address the aforementioned issues to some extent ^24,25,35,36^, however, the need for a deluge of training datasets poses another challenge in reconstruction efficiency and accuracy. Another emerging solution such as Fourier light field microscopy ^37–43^, addresses artifacts on the focal plane and provides a more uniform resolution by shifting the position of the MLA from the native image plane to the Fourier plane. However, it still faces challenges with limited resolving power for dense signals, a restricted depth of field, and a constrained fluorescence photon budget.

Zebrafish have proven to be a productive model for studying cardiovascular system development and related diseases. However, limitations in imaging and analysis techniques present challenges for studying the entire contracting heart in live models at cellular resolution. Our approach offers an efficient solution to investigate the cardiac structure and contractile function in zebrafish larvae, allowing us to track both periodic and arrhythmic dynamics across the entire heart. However, due to photon scattering and absorption, shallow photon penetration restricts the application of this strategy to zebrafish embryos and larvae, potentially limiting its use in adult zebrafish, which are optically opaque. While Casper ^44^ and Absolute fish lines ^45^ provide better transparency and reduced pigmentation, the enthusiasm for imaging applications is tempered by the challenges of additional nursery care, the need for double homozygous recessive propagation, and limited models for cardiovascular studies. In this context, the advancement of mutant, imaging techniques, or tissue-clearing approaches would significantly facilitate the development of cardiovascular study in live adult zebrafish.

## STAR★Methods

### Key resources table

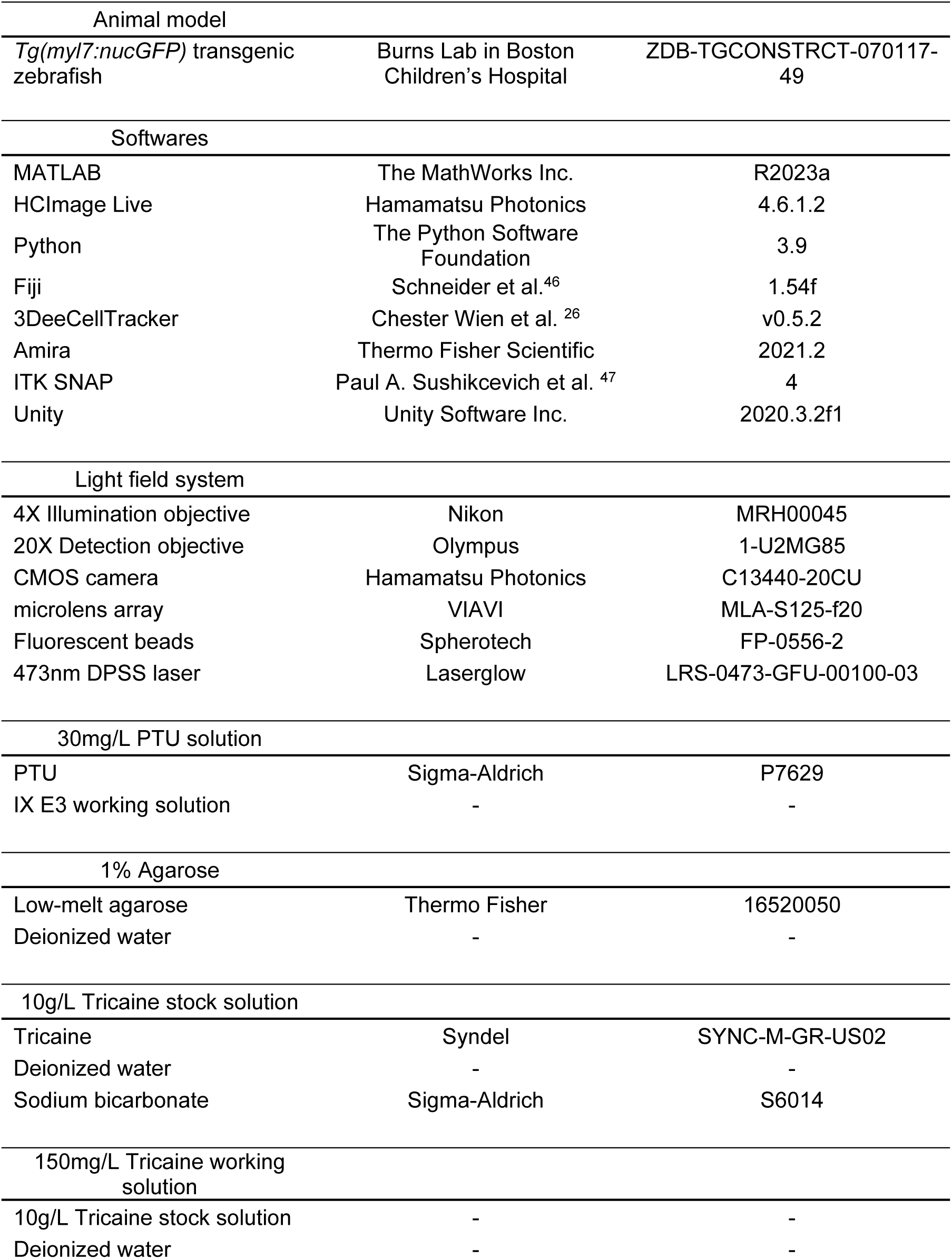
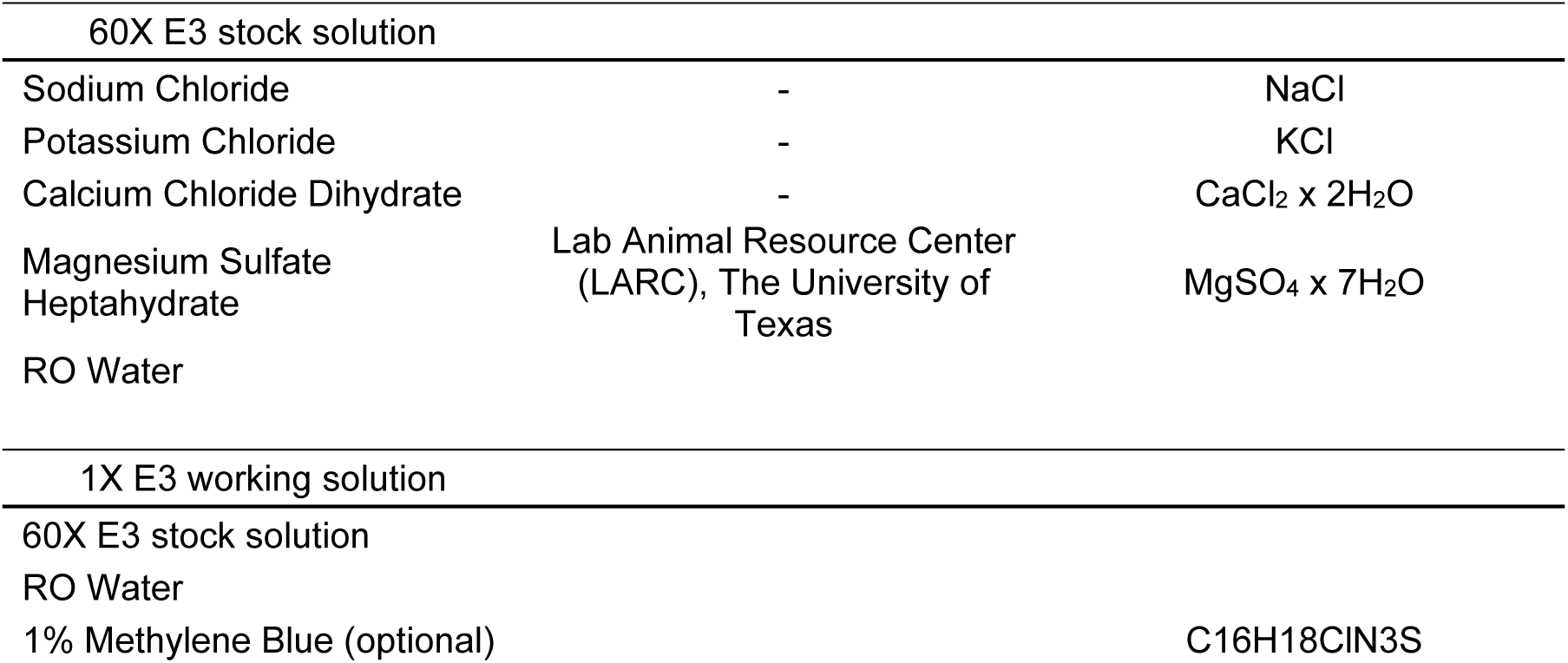

### Resource availability

#### Lead contact

Further information and resource requests should be directed to and will be fulfilled by the lead contact, Yichen Ding.

#### Materials availability

This study did not generate new unique reagents.

#### Data and code availability

All data and codes are available from the corresponding author upon reasonable request.

### Experimental model and subject details

#### Light field microscopy system

We built an in-house LFM imaging system using a continuous wave diode-pumped solid-state (DPSS) laser with a 473 nm wavelength (LRS-0473-GFU-00100-03, Laserglow Technologies) for illumination. A 5X beam expander (GBE05-A, Thorlabs) widened the laser beam, and a variable iris (ID25, Thorlabs) adjusted it to the preferred size. An achromatic doublet with a 125 µm focal length (AC254-125-A) focused the light on the back focal plane of an objective lens (Plan Fluor, 4×/0.13, Nikon), creating selective volumetric illumination across the entire zebrafish heart in the 3D-printed chamber. We collected the emitted fluorescence from the heart using a water-immersion objective lens (Plan Fluor, 20×/0.5, Olympus), the tube lens (ITL180, Thorlabs), an emission filter (FF01-441/511/717-25, Semrock), a microlens array (MLA-S125-f20, VIAVI), a relay lens (AF Micro-NIKKOR 60mm *f/*2.8D Lens, Nikon), and an sCMOS camera (Flash 4.0 v3, Hamamatsu). To match the NA of the MLA with that of the tube lens, we followed *N_MLA_* = *M/2NA_DL_* and selected the closest commercially available MLA for this project, where M is the magnification of the detection objective lens. Additionally, an LED was installed behind the sample holder to assist in identifying the position and orientation of the zebrafish during sampling mounting (**Supplementary Fig. S2**).

#### Simulation data

To analyze light propagation through the light field detection system, we utilized the Ray Optics Simulator ^48^ for tracing the light rays under various conditions. In addition, to simulate the PSF across varying depths, we developed a PSF generator in MATLAB, which allowed us to verify system alignment and compare the experimental results with theoretical analysis.

#### Preparation of the transgenic zebrafish sample

The transgenic *Tg(myl7:nucGFP)* zebrafish larvae from 3 to 6 dpf were used in this study. All animal protocols, experiments, and housing described in this manuscript were approved by the Institutional Animal Care and Use Committee (IACUC #20-07) of the University of Texas at Dallas. Zebrafish were maintained in an incubator at 28°C. To preserve optical transparency, we added 0.003% phenylthiourea (PTU) to the medium at 20 hours post-fertilization (hpf) to suppress pigmentation. Zebrafish larvae were anesthetized in 0.05% tricaine for 10 minutes for immobilization, then immersed in 0.5% low-melt agarose at 37°C within an FEP tube (refractive index: ∼1.33). The whole specimen, along with the FEP tube, was held by rotating stages to fine-tune the orientation of zebrafish heart after it was placed inside a chamber filled with DI water.

#### Image acquisition and reconstruction

We captured contracting hearts in live zebrafish larvae at different speeds, ranging from 30 to 200 frames per second (fps), depending on the varying fluorescence intensity in different fish. The reconstruction post data acquisition includes three main steps. First, we subtracted the average background signal intensity from the entire image in Fiji (version 1.54f). Second, we rectified the image by defining the pixel arrangement of the lenslets on the image, considering an area of 15*15 pixels behind each lenslet. To accurately apply this mask, we used a collimated beam to capture the exact positions of the lenslets, and incorporated it into reconstruction (**Supplementary Fig. S3)**. We employed 3D RL and EMS deconvolution methods to qualitatively and quantitatively compare their performance in cardiac images. A forward model was created to provide the initial estimation based on the theoretical PSF with system configurations (M = 20, NA_OL_ = 0.5, lenslet pitch = 125 µm, MLA focal length = 3125 µm), followed by comparison with the raw data. Based on the calculated error, an inverse model using the transposed PSF was applied to update the estimation ^18,19,22^. This process was iterated 15 times until convergence was achieved, enabling effective visualization of cardiomyocytes.

#### Cell displacement and velocity analysis

We utilized 3DeeCellTracker ^26^ to segment cardiomyocyte nuclei throughout a cardiac cycle. To enhance accuracy, we applied the watershed method ^49^ along with manual annotation. Subsequently, we performed 3D tracking using a feedforward network to identify cell positions across up to five cardiac cycles, with refinement achieved through PR-GLS ^50^. Tracking validation was performed by comparing predicted cell locations with the raw data.

#### VR analysis for assessment of cardiac activities

Our VR analysis framework enables a detailed study of zebrafish heart cells by processing and segmenting cardiac images to create a surface mesh for each individual cardiomyocyte nucleus. This framework allows for an intuitive analysis of cell contractility, morphology, and movement patterns throughout each cardiac cycle ^27–30^. To interact with a 4D heart model in VR, we started by obtaining cell tracking data from 3DeeCellTracker, then validated it to ensure consistent image intensity across all volumes, processing this step in Python. We transformed the heart models, which included all cells at various time points, into *.obj* and *.mtl* file formats and imported them into Unity. From there, we developed C# programs to enable 4D visualization and interactive analysis in VR. This VR platform supports cell selection, time point selection, pause functions, and lighting adjustments. Users can choose two cells to analyze their movement trajectories, velocities, volume, surface area, and relative distances over time. Additionally, the time point selection feature allows users to focus on specific points within the 4D data for deeper analysis. To highlight differences in cell displacement and speed, we applied a color-coding system where faster cells with greater overall movement appear brighter than others, making identifying cells with higher activity easier.

#### Data visualization

We used the “Fire” pseudo color in Fiji to indicate the fluorescence intensity distribution in 2D slices and employed the Z-stack Depth Color Code 0.0.2 plugin to encode the depth information in MIP images. To assess the performance of RL and EMS methods, we used logarithmic FFT to transform the spatial reconstructions to 2D frequency components. To enhance the analytic efficiency, we customized a Python script for tracking myocardial boundary contractions. The threshold-based detection allows for precise tracing of radial planar movements in any chosen direction.

#### Computational power

A workstation with an Intel(R) Xeon(R) W-3365 CPU, 256 GB of RAM, and an NVIDIA GeForce RTX 4090 GPU was utilized for the parallel computation, image post-processing, data analysis, and rendering.

## Acknowledgments

We express our gratitude to Dr. Caroline Burns’s group at Boston Children’s Hospital for generously sharing the transgenic zebrafish. We thank Ms. Elizabeth Ibanez and Mr. Adam Lavitz for their help in husbanding zebrafish at UT Dallas. We appreciate all the constructive comments provided by D-incubator members at UT Dallas.

## Author contributions

AS: Optical system design, Methodology, Zebrafish handling, Software, Validation, Formal Analysis, Investigation, Writing – Original Draft, Writing – Review & Editing, Visualization. RP, KC: Zebrafish handling, Software, Validation, Formal Analysis, Investigation, Writing – Original Draft, Writing – Review & Editing, Visualization. MA, XZ: Optical system design, Validation, Writing – Original Draft, Writing – Review & Editing. SSH, JB, JC, JL, BF, and JY: Software, Validation, Writing – Original Draft, Writing – Review & Editing. YD: Optical system design, Methodology, Zebrafish handling, Software, Validation, Formal Analysis, Investigation, Writing – Original Draft, Writing – Review & Editing, Visualization, Supervision, Project Administration, Funding Acquisition.

## Declaration of interest

There are no conflicts of interest to disclose.

## Funding

This work was supported by NIH R00HL148493 (Y.D.), R01HL162635 (Y.D.), NSF 2326628 (Y.D.), and UT Dallas STARS program (Y.D.).

## Notes

### Competing Interest Statement

The authors have declared no competing interest.

